# *Goldfinder*: Unraveling Networks of Gene Co-occurrence and Avoidance in Bacterial Pangenomes

**DOI:** 10.1101/2024.04.29.591652

**Authors:** Athina Gavriilidou, Emilian Paulitz, Christian Resl, Nadine Ziemert, Anne Kupczok, Franz Baumdicker

## Abstract

The pangenome is the set of all genes present in a prokaryotic species. Most pangenomes contain many accessory genes that are present in only some of the species members. Genes need to function together, and it has been suggested that selection for certain gene combinations affects the structure of prokaryotic pangenomes. Nevertheless, genes might also co-occur simply due to being linked on the genome, and efficient tools are needed to distinguish linkage from co-selection. Here we present Goldfinder, an approach to infer co-occurrence and co-avoidance between gene pairs by taking the phylogenetic relationships of the species into account. The approach is implemented in an efficient Python script available at https://github.com/fbaumdicker/goldfinder. We also provide scripts for clustering co-occurring genes and for visualizing the resulting co-occurrence and co-avoidance networks in Cytoscape. In comparison to the co-occurrence inference tool Coinfinder, Goldfinder finds fewer co-occurring pairs in a real species pangenome, suggesting that fewer spurious associations due to phylogenetic dependencies are detected. We conclude that Goldfinder is a fast and accurate tool to infer gene co-occurrence and co-avoidance, which will enable large-scale analyses to infer co-selected genes across bacterial pangenomes.

## Introduction

The pangenome of a population of bacteria or archaea is the set of all genes present in that population. Generally, each individual genome contains only a subset of the genes present in the pangenome (1), where the genes in a particular genome must work well together to form a functional organism. It has been observed that the fitness effect of a gene can depend on the genetic background (2, 3), suggesting epistatic effects between accessory genes in a pangenome. There are well-known examples of genes that need to function together to provide a benefit to the cell. For example, bacterial defense systems against mobile elements, such as CRISPR/Cas or restriction/modification, are often composed of multiple genes, where even additional interactions between defense systems have been proposed (4). Another well-known example are biosynthetic gene clusters, where different genes of a biosynthetic pathway must co-occur so that the complete pathway can be expressed and the relevant metabolite is produced (5). Similarly, some gene combinations might be unpreferred in a genome because they have redundant functions, or they even interact in a detrimental manner and lower the fitness of the cell.

Here, we aim to predict whether different combinations of genes are under selection. Based on the presence-absence patterns of accessory genes across strains, we infer whether genes tend to co-occur with each other or tend to avoid each other. However, gene presences might be associated or dissociated solely due to vertical inheritance: If two genes were gained in the same ancestral lineage, they show strong co-occurrence but are not necessarily functionally related. We will refer to this phenomenon as linkage by descent. In contrast, if two genes were gained and lost multiple times, each time in the same lineage, independently or as part of a larger gain or loss, this strongly suggests that they need to function together. Thus, phylogenetic correlation must be taken into account to disentangle linkage by descent from correlated presences and absences due to selection.

Various approaches have been developed in existing tools for analyzing gene co-occurrences and accounting for phylogenetic correlation. However, these approaches generally employ complex likelihood models (6–9) and are typically designed to investigate associations among few genes only. These approaches are expected to have long run times when applied to the complete pangenome. Other models are more computationally efficient but include only indirect phylogenetic information by restricting the set of considered genes (10, 11).

A related field are genome-wide associations studies (GWAS) where associations between genotypes and a phenotype are of interest. While accounting for the genome-wide linkage in prokaryotes is crucial in bacterial GWAS (12), the dependency on the underlying phylogeny is even more impactful when we consider all pairwise gene co-occurrences.

Here, we present Goldfinder (<u>G</u>ene <u>O</u>ccurrence accounting for <u>L</u>inkage by <u>D</u>escent), an approach to infer co-occurrence and avoidance due to selection by accounting for linkage by descent. Goldfinder is an efficient method which is based on a non-parametric approach and involves simulations along a user-provided phylogenetic tree.

## Main

### Approach

Goldfinder is a nonparametric and simulation-based approach that aims to identify genes that co-occur together or that avoid each other. Genes on bacterial genomes might co-occur even when they are not functionally related, due to vertical co-inheritance or population structure. Genes that offer a stronger selective benefit when present together in the same genome are likely to show a higher frequency of co-occurrence and are consequently often even found together in structures like mobile genetic elements, facilitating their horizontal co-transfer. To infer the pairs that co-occur due to co-selection or co-transfer instead of co-inheritance, Goldfinder accounts for genome-wide linkage and aims to identify only those pairs of genes that co-occur or avoid each other more often than expected by chance under the given phylogenetic relationships or population structure.

Goldfinder’s approach is inspired by the approach of the phylogenetic genome-wide association study (GWAS) tool treeWAS by Collins and Didelot (12) that has been designed for genotype-phenotype associations. Especially for bacterial datasets, GWAS methods need to consider the genetic relationships due to the genome-wide linkage. treeWAS accounts for this by simulating artificial genotypes along the phylogeny. Goldfinder uses a similar approach to infer genotype-genotype associations and considers all pairs of accessory genes in the pangenome to reconstruct networks of associated and dissociated genes. To detect gene pairs that co-occur due to co-transfer or co-selection, Goldfinder infers a score for each possible gene pair and simulates the null distribution of this score from simulated gene presence-absence data. An overview of the Goldfinder pipeline is shown in Figure 1. The simulations take the provided phylogenetic tree into account to prevent the identification of co-inherited gene pairs. Goldfinder requires two inputs: (i) A rooted phylogenetic tree that indicates the relationships of all strains in Newick format. If an unrooted tree is provided, Goldfinder will set a root via midpoint rooting. (ii) The gene presence/absence (PA) patterns of each strain, provided as a simple gene PA matrix or by the output of the pangenome analysis tools panX (13), Roary (14), or panaroo (15).

**Fig. 1.**
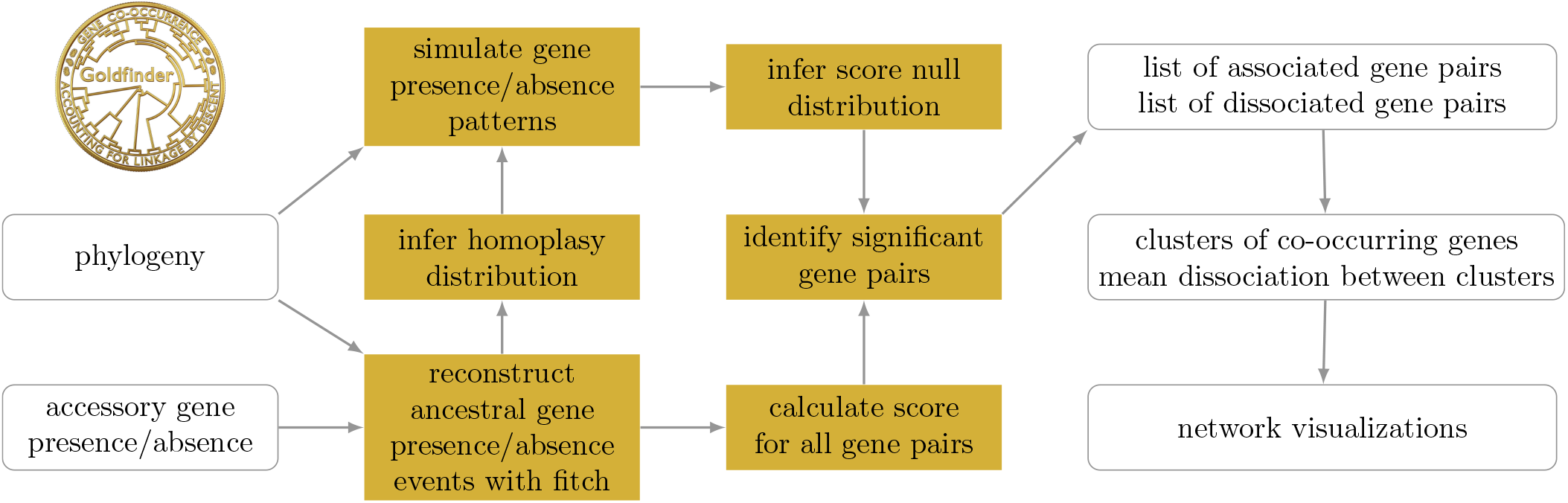
The Goldfinder workflow.

### Co-Occurrence and Avoidance Scores

Goldfinder infers a co-occurrence and an avoidance score for each possible gene pair. Different scores will differ in their ability to differentiate between co-occurrence resulting from genome-wide linkage or from selection. Goldfinder implements three different scores to indicate co-occurrence or avoidance of gene pairs: the score used in Coinfinder and the terminal and simultaneous scores as defined in treeWAS (12), where the terminal and the Coinfinder score are based only on the gene presences and absences in the terminal nodes and the simultaneous score is based on the reconstructed ancestral gene presences and absences at internal nodes of the phylogeny.

#### Ancestral gene presence-absence reconstruction

Based on the PA matrix and the phylogenetic tree, we reconstruct the presence or absence of genes at each ancestral node along the phylogeny using the Fitch algorithm (16). Whenever a switch of gene presence/absence is reconstructed, this indicates a gene gain or loss along the corresponding branch. See Figure 2. In addition to the calculation of the simultaneous score, the distribution of the number of reconstructed gene gains and losses forms the basis of our simulation approach.

**Fig. 2.**
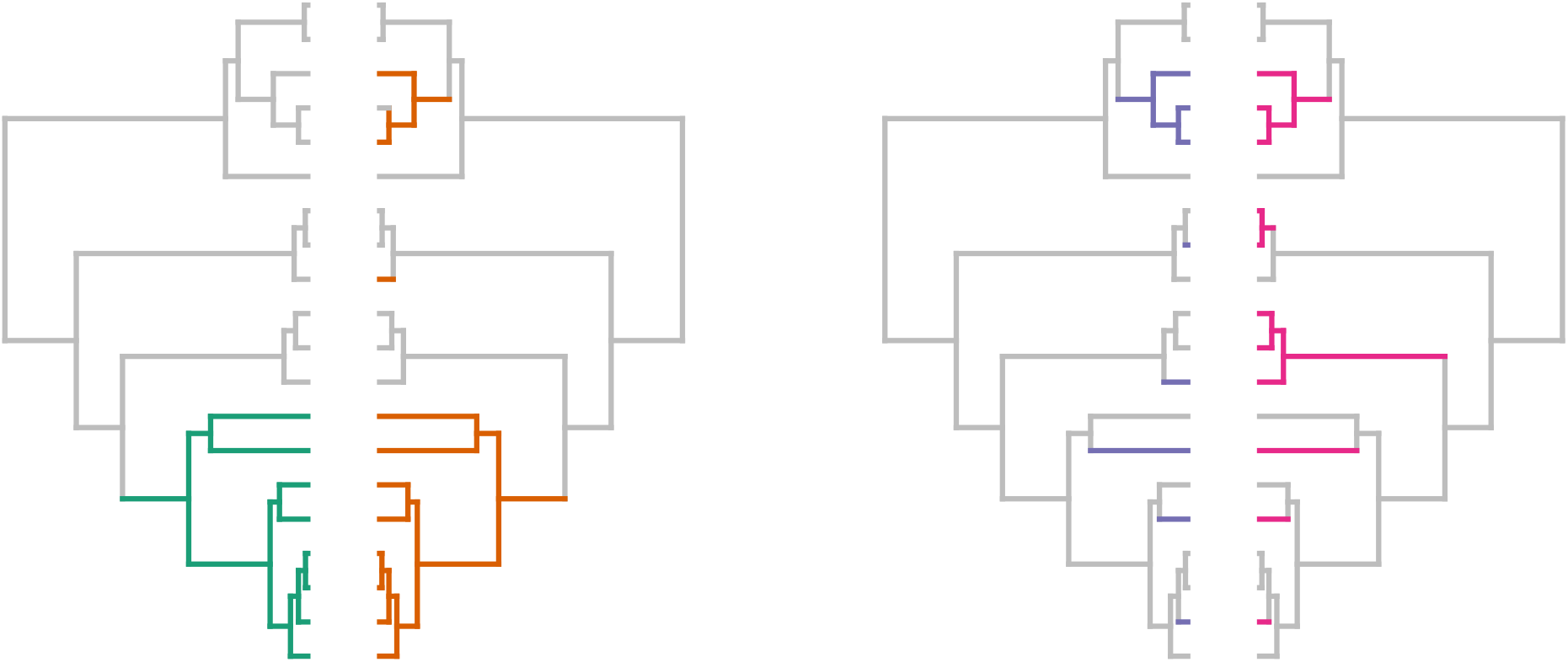
Four different gene presence-absence patterns and their reconstructed ancestral gain and loss events according to the Fitch algorithm. Along the colored lineages, the gene was reconstructed as present. The green gene has one reconstructed gene gain event. The orange gene has four state-changing events, three gene gains and one gene loss. The purple and the pink gene have six reconstructed gene gains each. The two gene pairs (green/orange and purple/pink) have the same terminal score (17) and coinfinder score (8) but different simultaneous scores (1 and 4). Thus, genes with the same score can have evolutionary paths with strikingly different probabilities, making it difficult to distinguish coinheritance from true associations or dissociations. Coinfinder tries to account for this by excluding genes with a low D-value such as the green gene here.

#### Score calculation

##### Terminal Score

For a pair of genes, the Terminal score of association *A*_*T*_ is given by *A*_*T*_ = *G*_*P*_ + *G*_*A*_ − *G*_*D*_, where *G*_*P*_ is the number of strains where both genes are present, *G*_*A*_ is the number of strains where both genes are absent, and *G*_*D*_ is the number of strains where one gene is present and the other is absent (i.e., they are different). The Terminal score for dissociation is *D*_*T*_ = −*A*_*T*_ . The scores are between −*n* and *n* (where *n* is the number of strains) and centered around 0, if the genes are randomly distributed.

##### Coinfinder Score

The Coinfinder score for association is *A*_*C*_ = *G*_*P*_ and the Coinfinder score for dissociation is *D*_*C*_ = *G*_*D*_. The scores are between 0 and *n*.

##### Simultaneous Score

The Simultaneous score of association is *A*_*S*_ = *B*_*S*_ − *B*_*D*_, where *B*_*S*_ is the number of branches in the tree where both genes are lost or both are gained and *B*_*D*_ is the number of branches where one gene is lost and one gene is gained. The Simultaneous score of dissociation is *D*_*S*_ = −*A*_*S*_. The scores vary between −2*n* + 2 and 2*n* − 2.

#### Score comparison

In the application scenario of treeWAS, to detect associations between genotypes and phenotypes, dependencies due to coinheritance are typically weak and consequently, the terminal score can provide useful insights. However, in the case of co-occurrence or avoidance between genes, the terminal score and the coinfinder score do not take into account the phylogenetic relationships between strains, which naturally generate a correlation between the evolutionary history of all genes. Therefore, both scores are not expected to effectively differentiate between gene pairs that co-occur because of a few gene gain or loss events that likely coincide along the same longer branches of the phylogeny and on the other side those gene pairs that co-occur as a result of many less likely coincident gene gain or loss events on shorter branches; see Figure 2. Thus, we expect that the simultaneous score is more suited to distinguish genome-wide linkage and selection and is therefore the recommended default score in Goldfinder.

### Simulation of gene presence-absence patterns and the resulting null distribution

Next, we aim to infer the score null distributions, where no association between genes due to function or co-inheritance occurs. To this end, we simulate gene presence-absence evolution along the underlying phylogeny, where all genes evolve independently. Thus, all co-occurrences arise solely due to co-inheritance in the simulations. To mimic the gene frequencies in the provided pangenome, the simulation relies on the distribution of the total number of reconstructed gains and losses per gene, which we refer to as the homoplasy distribution. Other authors have referred to this number as the exchangeability of a gene (7).

#### Homoplasy distribution

From the ancestral reconstructions, we infer the homoplasy distribution, i.e., the distribution of the number of state-changing events reconstructed by the Fitch algorithm (16). Here, this refers to the number of branches with a reconstructed gene gain or loss, as illustrated in Figure 2.

We noticed that the homoplasy distribution strongly depends on the ancestral state reconstructed at the root of the tree. Genes that are present at the root are often part of the extended core genome and thus have a high probability of having a low number of gene losses (and regains). In contrast, genes that are not present at the root are often mobile genes that are more frequently gained and lost again. Thus, we infer the homoplasy distribution conditional on the ancestral state at the root of the phylogenetic tree, which enables us to more accurately replicate the distribution of accessory genes in the pangenome.

#### Simulating gene presence-absence evolution

To simulate gene PA patterns, we first determine the ancestral root state according to the empirical probability of the reconstructed gene presence or absence at the root. Then the number of gene-presence-changing events is chosen according to the homoplasy distribution conditioned on the root state. Finally, a PA-pattern with that number of events is simulated, by distributing gene gains and losses among the branches of the phylogenetic tree with probabilities proportional to their branch length.

To treat simulated data and real data identically, we use the presence-absence patterns of the neutrally simulated genes to reconstruct ancestral states and calculate the scores between all gene pairs. These simulated scores constitute the null distribution of the co-occurrence scores.

Our approach assumes that the rate of gene gains and losses is constant for all branches of the phylogeny. Goldfinder is thus a tool tailored for the analysis of pangenomes of closely related strains of the same species, and applications to gene co-occurrence across different species should be considered with caution.

#### Refinement of the simultaneous score

To determine which gene pair co-occurrences are significant, we are mainly interested in the extreme values among the possible simultaneous scores. However, for the simultaneous score, those values are only reached rarely, since values above *k* can only be reached if the fitch score of both genes is at least *k*. Thus, we would have to simulate many PA patterns to get a high resolution of the tail of the score distribution. Instead, we simulate the score distribution in two steps. First, we simulate gene PA patterns drawing from the complete homoplasy distribution. In a second step, we refine the empirical distribution of the more extreme scores by simulating only the co-occurrences of genes with a high number of PA changes and we correct for their frequency in the pangenome. This property allows us to dramatically reduce the number of simulations required to obtain a precise estimate of extreme simultaneous scores without any loss in accuracy.

### Networks of Gene Co-occurrence and Avoidance

#### Identifying significant gene pairs and multiple testing corrections

Based on the null distribution, a p-value is determined for each gene pair as the proportion of simulated scores that exceed the score of the gene pair. Depending on the significance level, this results in a list of significantly associated or significantly dissociated gene pairs.

Goldfinder provides multiple options to address the multiple-testing problem that naturally arises when looking at all possible pairs of genes in the pangenome. Golfinder can output Bonferroni-corrected and false discovery rate (FDR)-based p-values, as well as the uncorrected p-values. The Bonferroni correction assumes that the tests are independent, which is clearly violated for pairs of co-occurrence patterns; thus the Bonferroni correction tends to exclude too many gene pairs. Therefore, we set an FDR-based correction as the default in Goldfinder.

#### Reconstructing clusters of co-occurring genes

All gene pairs with a significant co-occurrence can be connected, resulting in the formation of an extensive network of co-occurring genes. We observed that the largest connected component of this network can include almost all genes (see also Figure 3 (C)). However, some subsets of this network will exhibit dense clusters of co-occurring genes. To extract these, we employ the Markov Cluster Algorithm MCL (17) using the inferred p-values as input. The resulting distinct clusters are composed of genes that have low inferred p-values with each other and form highly interconnected groups of co-occurring genes (see also Figure 3 (D)). Note that some of the co-occurring gene pairs will connect different clusters of co-occurring genes as identified by the MCL algorithm. Naturally, such connections occur for only a few gene pairs between the identified clusters. In Figure 3 (C) The complete gene co-occurrence network is displayed in a way that genes belonging to the same MCL clusters are positioned closer to each other, enhancing the visual representation.

**Fig. 3.**
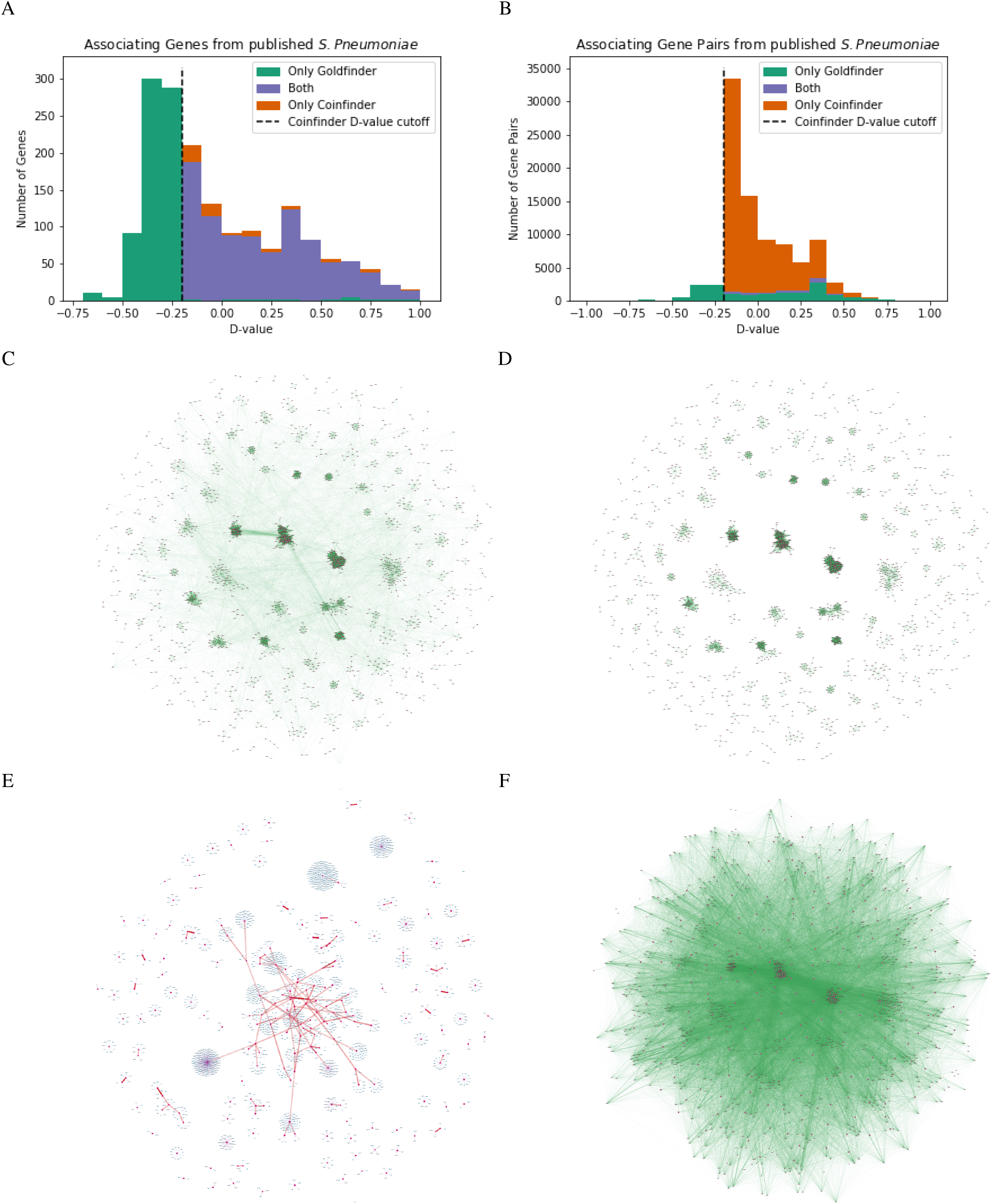
Comparision of Goldfinder and Coinfinder: **A:** The number of genes that are in at least one gene-gene association in Goldfinder, Coinfinder, or both, for different D-values. **B:** The number of gene-gene associations for different D-values, found by Goldfinder, Coinfinder, or both. **C:** A visualization of the network of associated gene pairs as identified by Goldfinder. The dots represent individual genes and if two genes co-occur according to Goldfinder they are connected by a green line. **D:** A visualization of the sub-networks of associated gene pairs as identified by the MCL tool based on the Goldfinder p-values. Only the connections within MCL clusters are shown. **E:** The visualization of the clusters of co-occurring genes and dissociations between those clusters as inferred by Goldfinder. Clusters of co-occurring genes are represented by the red centered nodes, blue nodes correspond to genes within the cluster. Dissociated clusters of co-occurring genes are connected by a red line, where the fraction of dissociated gene pairs determines the opacity of the line. **F:** A visualization of the network of associated gene pairs as identified by Coinfinder. The nodes represent individual genes and if two genes co-occur according to Coinfinder they are connected by a green line. The nodes of the genes are placed at the same positions as in C and D.

#### Mean avoidance between clusters of co-occurring genes

Naturally, avoiding genes cannot be as easily clustered as co-occurring genes. Instead, we consider the mean dissociations between clusters of co-occurring genes as follows: First, for all pairs of MCL clusters, we identify all genes that have a significant avoidance score with more than half of the genes of the other MCL cluster. The fraction of genes identified from both MCL clusters provides the mean dissociation between the clusters.

We visualize these dissociations between all clusters of co-occurring genes, where at least one gene avoids at least 50% of the other cluster (Figure 3 (E)).

#### Graphical visualization in Cytoscape

Goldfinder outputs a number of visualizations of the gene co-occurrence network that can be explored in the Cytoscape tool (18) with the help of the py4cytoscape Python package (19), implemented in a jupyter notebook (.ipynb) file, which is provided. The main visualization of the gene co-occurrence network is focused on MCL clusters and can omit or include the gene associations between different MCL clusters; see Figure 3 (C) and (D). In addition, we provide a visualization of the mean dissociation between clusters of co-occurring genes; see Figure 3 (E).

### Application to *Streptococcus pneumoniae*: Goldfinder identifies specifically the phylogeny-independent gene co-occurrences

We investigated how the explicit consideration of the underlying phylogeny in Goldfinder influences the detection of co-occurring gene pairs. In particular, we investigate the detected gene pairs in relation to how the genes correlate with the provided phylogeny. A strong correlation between the presence or absence of a gene and the phylogeny can be indicated by a low D-value (20). The D-value is also used in Coinfinder to exclude genes that closely follow the phylogeny from the co-occurrence analysis.

However, for genes with higher D-values and thus a low correlation between the presence and absence of genes and the phylogeny, the distribution of pairwise co-occurrence is still distorted by the shared phylogenetic relationship. Goldfinder thus uses a simulated null distribution conditionend on the provided phylogeny, whereas Coinfinder assumes a phylogeny-independent null distribution of gene presence and absence. Consequently, Goldfinder can (and has to) consider all genes, whereas Coinfinder excludes genes with a low D value, i.e., a strong correlation with the provided phylogeny, to partly compensate for the assumption of a phylogeny-independent null distribution. In addition, Goldfinder uses the simultaneous score based on the terminal and the reconstructed internal states of the phylogeny, while Coinfinder uses a score based solely on the terminal states. These significant differences to Coinfinder, enable Goldfinder to identify specifically the phylogeny independent gene co-occurrences.

To compare the co-occurrences found by Goldfinder and Coinfinder, we reanalyzed the *Streptococcus pneumoniae* pangenome data of Whelan et al. (10) using the default parameters of each pipeline and determining the D-value cut-off as recommended by Coinfinder.

When looking at the genes that are in at least one co-occurrence relation, Goldfinder and Coinfinder find nearly the same number of genes, except for genes below the D-value threshold (Figure 3 (A)). However, when looking at all the gene pairs identified, the number of associated gene pairs is much higher for Coinfinder and this difference increases for lower D-values. In total, Goldfinder identifies 18443 significant gene pairs wheras Coinfinder connects 76416 gene pairs.

## Conclusions

Here, we present Goldfinder, an approach to infer gene co-occurrence and avoidance due to co-selection or co-transfer. Goldfinder is an efficient method that can be applied to all accessory genes of a species, that is, its pangenome. By taking a phylogeny of the strains into account, which is integral in the score calculation itself and in the simulation of the neutral null distribution, Goldfinder is able to distinguish co-selection from pairs that co-occur or avoid each other due to genome-wide linkage. Pairs that are inferred to be under co-selection are clustered in a network that is visualized in an appealing way. In addition, dissociated gene pairs can also be estimated and displayed in a separate network. We are aware of only one other tool to infer co-selection that is also designed for pangenomes, Coinfinder. The comparison between both tools shows that Goldfinder infers fewer gene pairs under co-selection but more total genes involved in co-selection, suggesting that more specific gene pairs are inferred. The availability of a fast and accurate tool to infer gene co-occurrence and avoidance will enable large-scale analyses to infer co-selected genes across whole species pangenomes. Goldfinder will enhance our understanding of the evolutionary forces that shape prokaryotic pangenomes.

## DATA AND CODE AVAILABILTIY

The software Goldfinder is available at https://github.com/fbaumdicker/goldfinder.

## ACKNOWLEDGEMENTS

AG thanks for the support of the Deutsche Forschungsgemeinschaft (DFG; Project ID No. 398967434-TRR 261). NZ is supported by the German Center for Infection Research (DZIF) (TTU 09.716). NZ and AG thank CMFI (Germany’s Excellence Strategy–EXC 2124–390838134) for structural support. FB is funded by the Deutsche Forschungsgemeinschaft (DFG, German Research Foundation) under Germany’s Excellence Strategy – EXC number 2064/1 – Project number 390727645, and EXC 2124 – Project number 390838134.

